# Engineering of soluble bacteriorhodopsin

**DOI:** 10.1101/2024.11.20.624543

**Authors:** Andrey Nikolaev, Yaroslav Orlov, Fedor Tsybrov, Elizaveta Kuznetsova, Pavel Shishkin, Alexander Kuzmin, Anatoly Mikhailov, Yulia S. Galochkina, Arina Anuchina, Igor Chizhov, Oleg Semenov, Ivan Kapranov, Valentin Borshchevskiy, Alina Remeeva, Ivan Gushchin

## Abstract

Bacteriorhodopsin is a seven-helical light-driven proton pump and a model membrane protein. Here, we report engineering of soluble analogues of bacteriorhodopsin, NeuroBRs, which bind retinal and photocycle under illumination. We also report the crystallographic structure of NeuroBR_A, determined at anisotropic resolution reaching 1.76 Å, that reveals a conserved chromophore binding pocket and tertiary structure. Our results highlight the power of modern protein engineering approaches and pave the way towards wider development of molecular tools derived from membrane proteins.

---

Membrane proteins are notoriously difficult objects for study. Consequently, significant effort was applied to develop efficient solubilization methods^1,2^. Alternatively, several protein engineering approaches were devised to engineer soluble analogues of membrane proteins^2^. In two cases (KcsA and nAChR), successful design of soluble variants was reaffirmed by obtaining NMR structures^3–5^. Later, QTY methodology was introduced, which, in particular, allowed generation of soluble forms of GPCRs able to bind their expected ligands^6^, although QTY proteins have so far resisted structure determination efforts^2^. More recently, a plethora of easy-to-use machine learning-based protein engineering techniques were developed displaying high success rates. ProteinMPNN, utilizing message-passing neural networks, predicts amino acid sequences that would fold into desired shape, taking the protein backbone coordinates as an input^7^. Proteins engineered with ProteinMPNN express in greater amounts, are more soluble and stable and, in some cases, have higher activity^8^. Recently, modified ProteinMPNN was used to generate soluble proteins with folds closely matching those of claudin, rhomboid protease and GPCRs, as confirmed using X-ray crystallography^9^. Whereas the interior of these proteins was completely redesigned, grafting of functional residues on the protein surface endowed them with the ability to bind natural partners of their prototypes^9^.

Bacteriorhodopsin (BR), a protein from the archaeon *Halobacterium salinarum* discovered in 1971^10^, is the prototypical member of the microbial rhodopsin family, and probably the best studied membrane protein overall^11^. It consists of seven α-helices (labeled A to G) and covalently binds cofactor retinal, which undergoes isomerisation upon illumination. Photoactive proteins from this family are ubiquitous in the biosphere^12^, and are being used extensively in optogenetics^13,14^. Several unsuccessful attempts to engineer soluble bacteriorhodopsin were previously reported^2^.

Here, we applied SolubleMPNN^9^ to engineer soluble variants of bacteriorhodopsin. In line with previous work applying neural networks to ligand-binding proteins^8,15,16^, we fixed the retinal-binding pocket residues (Fig. 1a), while allowing all other residues to be mutated. We generated 52 sequences using SolubleMPNN and used ColabFold^17^ to select three sequences, dubbed NeuroBR_A, B, and C, for which general folds and retinal binding pockets were confidently predicted to be close to those of the wild type protein (Fig. 1b). Sequence identities between WT BR and NeuroBR_A, B, and C are 41.2, 43.4, and 44.7 %, respectively. We then conducted molecular dynamics simulations of the three designs, which revealed potential stability of the developed variants (Fig. 1c). Phylogenetic analysis places NeuroBRs in the same branch as WT BR, but much further from the origin (Fig. 1d). While the exterior of the 7-helical bundle is mostly hydrophobic in the WT BR, predicted structures show a lot of polar or potentially charged residues on the protein surface (Fig 1e).

**Figure 1.**
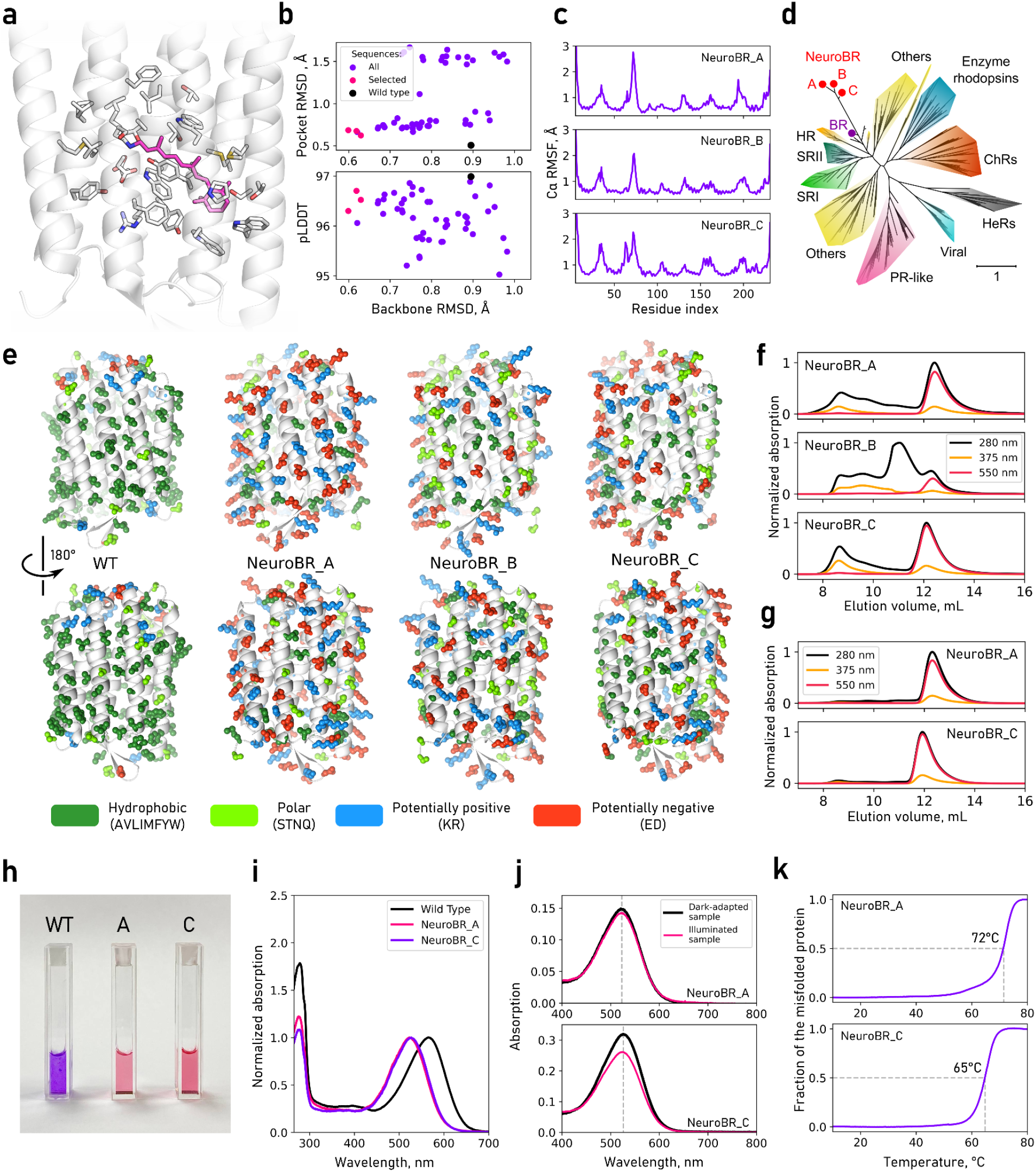
Sequence design and initial characterisation of soluble BR variants. **a**, Chromophore-binding pocket in WT BR. **b**, Backbone RMSD, pocket RMSD and pLDDT of AlphaFold-predicted models for WT BR and SolubleMPNN-generated sequences. **c**, Amplitude of thermal fluctuations of C_α_ atoms of engineered BR variants in MD simulations. **d**, Phylogenetic tree for NeuroBRs and representative microbial rhodopsins (based on dataset by Rozenberg *et al*.^12^). **e**, Surface amino acid properties for WT and engineered BR variants. Histidines and cysteines are absent from WT and engineered variants, except for 6×His tags added to NeuroBRs for metal affinity purification (not shown). **f**, SEC profiles for NeuroBRs reconstituted with retinal. **g**, SEC profiles for monomeric fractions of NeuroBR_A and C after one week incubation at 4 °C. **h**, Visual appearance of WT BR in purple membranes and NeuroBR_A and C in detergent-free buffer. **i**, Absorbance spectra of WT and engineered BR. **j** Comparison of absorption spectra of dark-adapted and illuminated samples of NeuroBR_A and C, **k**, Thermal denaturation of engineered BR variants.

Next, we proceeded with experimental characterization of NeuroBRs. Usually, during heterologous expression of retinal-binding proteins in *Escherichia coli*, the retinal is provided exogenously as it is not synthesized by the bacterium. Despite retinal supplementation, all three proteins were expressed and purified (Fig. S1) in colorless apo-forms. We believe this to be the consequence of the cell wall being impermeable to the hydrophobic retinal, originally added as an ethanol solution. Consequently, we incubated the purified proteins in yellow-colored retinal solution overnight and observed formation of pink-colored protein complexes. Size exclusion chromatography (SEC) revealed that after the incubation the holo-proteins were present as mixtures of the proteins in oligomeric orange and monomeric pink forms (Fig. 1f). Separated pink form remained stable and monomeric for at least a week (Fig. 1g,h). NeuroBR_B was discarded from further study as its pink form was unstable, converting with time into an orange form (see Supplementary Text for the details).

Absorption maxima of purified monomeric NeuroBR_A and C were observed at 523-526 nm (Fig. 1i), shifted from 548-568 nm (depending on conditions) observed for WT BR. Whereas WT BR displays dark adaptation (blue shift of the absorption maximum), no alterations in NeuroBR spectra were observed after storing in darkness for 12 hours (Fig. 1j). The proteins were stable with thermally induced transitions to the orange form observed at 72 and 65 °C (Fig. 1k). Finally, the proteins were observed to undergo a photocycle (Figs. 2,S2,S3). Three distinct intermediates were clearly resolved: early red-shifted K_535_ intermediate, strongly blue-shifted M_400_ intermediate (presumably corresponding to the deprotonated form of the Schiff base), and late red-shifted O_585_ intermediate. Decay rates of the intermediates of NeuroBR_C, and the K state of NeuroBR_A were essentially independent of pH, whereas decay rate of the M and O intermediates increased with increasing proton concentration, indicating direct proton uptake from the solution during the M state decay. Above pH 8.0, NeuroBR_A converted into a long-lived orange state upon illumination, partially recovering to the pink form over several hours in darkness. Slowdown of the photocycle of NeuroBR variants compared to the WT protein probably follows from the lack of the proton donor residue, as in the D96N BR variant^18,19^.

**Figure 2.**
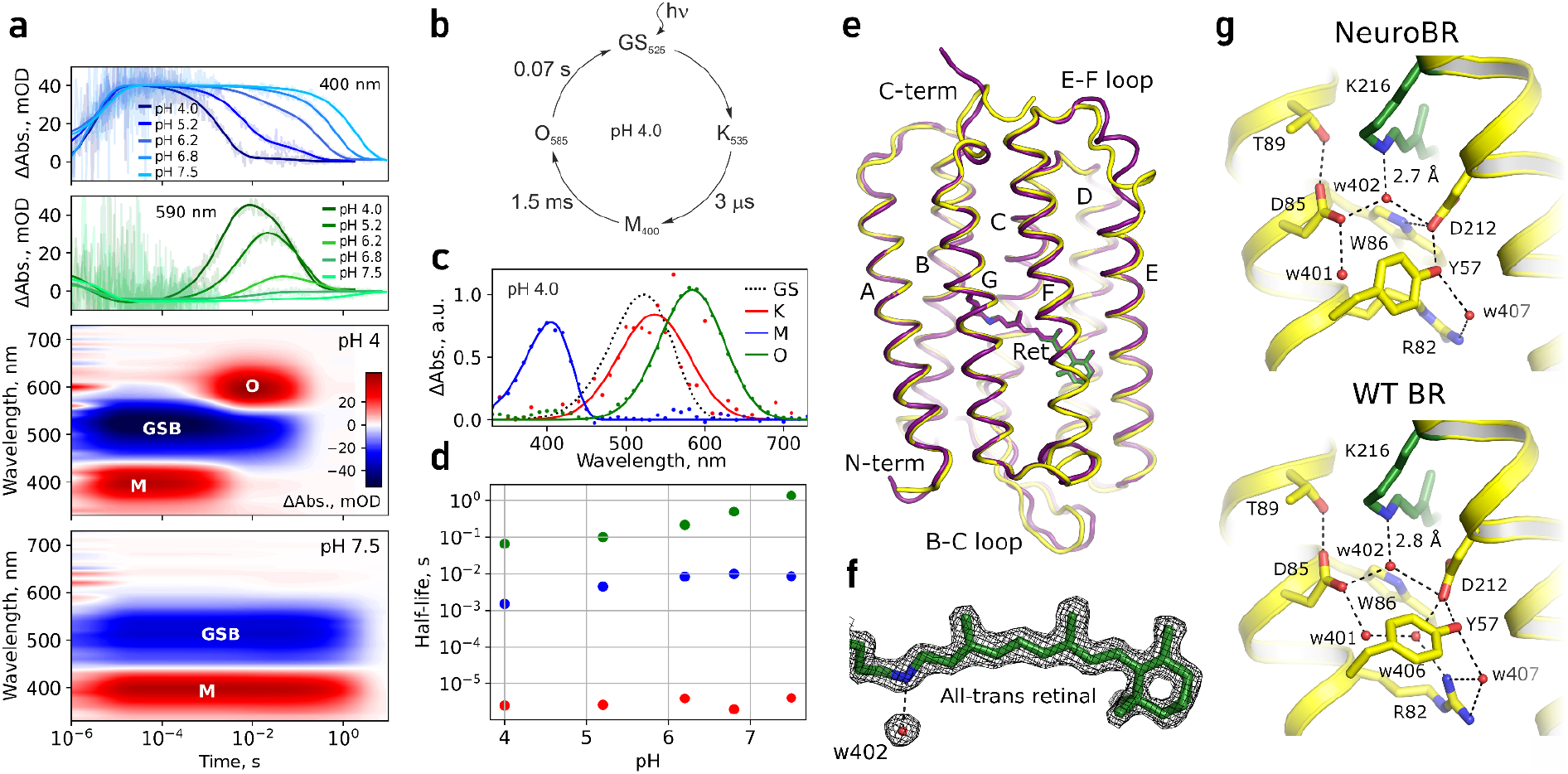
Photocycle and structure of soluble BR variant NeuroBR_A. **a**, Changes in absorbance of NeuroBR_A in solution after flash illumination. M and O correspond to areas where absorbance raises due to formation of putative M and O photocycle intermediates. GSB is the ground state bleaching area where absorbance corresponding to the ground state is diminished due to formation of photocycle intermediates. **b**, Model of NeuroBR_A photocycle at pH 4.0, where the different photocycle intermediates are clearly distinguishable. **c**, Recovered absorption spectra of NeuroBR_A photocycle intermediates. **d**, Dependence of NeuroBR_A photocycle intermediate half lifes on pH. **e**, Overlay of backbone and retinal structures for WT BR (purple) and NeuroBR_A (yellow and green). **f**, Weighted 2F_o_-F_c_ electron densities around the retinal contoured at 2 σ. **g**, Comparison of retinal Schiff base environment in WT BR and NeuroBR_A. w406 is absent in NeuroBR_A and R82 changes its conformation.

Following photophysical characterization, we attempted crystallization of NeuroBR_A and NeuroBR_C. We observed the formation of box-shaped crystals of NeuroBR_A in 2 days (Fig. S4). The crystals diffracted anisotropically, with resolution cut-offs for the best single crystal dataset of 1.76/1.78/2.21 Å. The structure was solved using molecular replacement with an AlphaFold-generated model, with 4 molecules in the asymmetric unit. No apparent oligomerization interfaces were observed; the proteins pack in layers in the crystal (Fig. S5).

Whereas NeuroBR sequences correspond to amino acids 5-231 of WT BR, below we use WT BR numbering of amino acids for NeuroBR for clarity and for correspondence with earlier literature. Electron density allowed clear assignment of most amino acids, with the exception of flexible N- and C-termini and the amino acid 161 in the chain D; this amino acid is located in the E-F loop that is also disordered in many WT BR structures. The overall fold is extremely well conserved (Fig. 2e) with the root mean square deviation (RMSD) of all heavy atom positions of 1.43-1.55 Å and retinal-binding pocket atom positions of 0.69-0.77 Å. In all four molecules, the retinal is in apparent all-trans conformation (slightly more straightened compared to the WT structure), and the conserved water molecule W402 is clearly resolved near the retinal Schiff base (Fig. 2f). Interestingly, Arg82 is reoriented away from the Schiff base compared to its position in the ground state of WT BR (Fig. 2g). The only notable deviations of the obtained structure from the design are 2.7 Å displacement of the beta-hairpin element (B-C loop) and rearrangement of the E-F loop (disordered in most WT BR structures; Fig. 2e).

Engineering of NeuroBR_A presents a clear-cut and unambiguous example of converting a membrane protein into a soluble one, while retaining the ligand-binding ability of its intramembrane part and its core function. This soluble bacteriorhodopsin analogue may be used for probing the retinal photophysics and photochemistry in a unique environment of α-helical soluble protein, complementing as a model system the soluble rhodopsin mimic based on the human cellular retinol binding protein II^20^. Similar methodology may probably be used to generate soluble analogues of drug target membrane proteins for easier screening of potential hit molecules. Overall, our results further highlight the power of modern protein engineering approaches and pave the way towards wider development of molecular tools derived from membrane proteins.

## Materials and Methods

### Computational protein engineering

To preserve the natural ligand-binding ability of BR, we fixed the binding site residues during re-engineering^8,15,16^. Using the high resolution structure of ground state BR (PDB ID 7Z09^21^) as a reference, we fixed residues with side chain atoms within 5 Å of the retinal and glycines with heavy atoms within the same distance of the retinal. We also fixed the identity of Y57 and R82 for proper coordination of D212, and fixed T178 for proper coordination of W182. Finally, we fixed the region of the G helix containing residues 212-220 around the retinal-binding K216. In total, 34 residues were fixed: 20, 49, 50, 53, 57, 82, 83, 85, 86, 89, 90, 93, 118, 119, 122, 138, 141, 142, 145, 178, 182, 185, 186, 189, 208, 212, 213, 214, 215, 216, 217, 218, 219, 220.

We applied ProteinMPNN^7^ retrained only on structures of soluble proteins^9^ to coordinates of the residues 5-231 of WT BR structure. We chose a SolubleMPNN model that predicts amino acid identity considering 48 neighboring amino acids and was trained using Gaussian noise with SD of 0.2 Å, with sampling temperature of 0.1. In total, we generated 52 sequences and predicted the corresponding atomistic models using AlphaFold2-based ColabFold v. 1.5.5 with alphafold2_ptm model^17,22^, with the number of recycles set to 6. For each sequence, 5 models were obtained. Best model for consequent analysis was selected using average pLDDT. No AMBER relaxation was performed afterwards. Next, three sequences were selected for further characterization using ColabFold-predicted metrics: average pLDDT, root mean squared deviation (RMSD) of all C_α_ atoms and RMSD of heavy atoms of fixed residues. The sequences were dubbed A, B, and C in descending order of average pLDDT. Metal affinity tags consisting of six histidines were added via a GSG linker at the C-terminus of each sequence. Final amino acid and nucleotide sequences are available as Data S1 and S2. Properties of respective proteins are listed in Table S1.

### Molecular dynamics simulations

NeuroBR_A, B, C were simulated in the monomeric form. Initial models were obtained by combining the previously obtained models and positions of internal water molecules and retinal from the crystallographic structure of WT BR in the ground state (PDB ID 7Z09) using PyMOL^23^. Dodecahedron unit cell contained 12455 water molecules, 50 Na^+^ and 38 Cl^-^ ions for the NeuroBR_A; 13367 water molecules, 48 Na^+^ and 40 Cl^-^ ions for the NeuroBR_B; 12472 water molecules, 54 Na^+^ and 38 Cl^-^ ions for the NeuroBR_C. All atomistic systems were prepared using the tools of GROMACS 2022^24^. Protonation states of all titratable residues were assigned in accordance with pH of 7.0 using PROPKA 3.5.1^25^; the retinal Schiff base was protonated.

MD simulations were conducted using GROMACS 2022, with CHARMM36 force field^26^, and TIP3P water model^27^. Parameters for the retinal bound to lysine were adapted from ref.^28^. Systems were energy minimized using the steepest descent method, thermalized, equilibrated and simulated for 200 ns using the leapfrog integrator with a time step of 2 fs, at a reference temperature of 303.15 K and at a reference pressure of 1 bar. Temperature was coupled using Nosé-Hoover thermostat^29^ with coupling constant of 1 ps^−1^. Pressure was coupled with isotropic Parrinello-Rahman barostat^30^ with relaxation time of 5 ps and compressibility of 4.5⋅10^−5^ bar^−1^. Coordinates of the systems were collected every 20 ps.

The simulations were performed using periodic boundary conditions. The covalent bonds to hydrogens were constrained using the LINCS algorithm^31^. The nonbonded pair list was updated every 20 steps with the cutoff of 1.2 nm. Force based switching function with the switching range of 1.0–1.2 nm and particle mesh Ewald (PME) method^32^ with 0.12 nm Fourier grid spacing and 1.2 nm cutoff were used for treatment of the van der Waals and electrostatics interactions. The Python library MDAnalysis^33^ was used for analyses.

### Phylogenetic analysis

We used MAFFT 7.520^34^ to align sequences of designed proteins NeuroBR_A, B, and C to the trimmed alignment of 315 sequences from the review by Rozenberg *et al*.^12^ as available via GitHub (https://github.com/BejaLab/RhodopsinsReview/blob/main/data/). Phylogenetic tree was constructed using IQ-TREE 1.6.10^35^ with additional options that correspond to those used by Rozenberg *et al*.^12^ (-m TEST -bb 1000 -pers 0.2 -nstop 500). Tree image was generated using TreeViewer 2.2.0^36^.

### Protein expression and purification

Protein-coding sequences were synthesized *de novo* and cloned into pET-28a(+) plasmid via XbaI (TCTAGA) and BamHI (GGATCC) restriction sites. *E. coli* strain C41 (DE3) cells transformed with protein-encoding plasmids pET-28a(+) were cultured in shaking flasks in 400 ml TBP-5052, one liter of which was prepared by mixing 930 ml of TB (12 g/L tryptone, 24 g/L yeast extract), 1 ml of 1 M MgSO_4_ solution, 20 ml of 50×5052 (250 g/L glycerol, 25 g/L glucose, 100 g/L α-lactose) and finally 50 ml of 20xNPS (66 g/L (NH_4_)_2_SO_4_, 136 g/L NaH_2_PO_4_, 142 g/L Na_2_HPO_4_). Kanamycin was added to the concentration of 50 mg/L. Cell cultures were incubated at 37 °C until reaching optical density of 0.5-0.7. Protein expression was induced by addition of 1 mM IPTG and followed by incubation for 20 h at 20 °C. Harvested cells were resuspended in lysis buffer containing 300 mM NaCl and 50 mM Tris-HCl, pH 8.0 and were disrupted in an M-110P Lab Homogenizer (Microfluidics, USA). Cell membrane fraction was removed from the lysate by ultracentrifugation at 100 kg for 45 minutes at 10 °C. Clarified supernatants were incubated overnight with Ni-nitrilotriacetic acid (Ni-NTA) resin (Qiagen, Germany) with constant stirring at 4 °C. Supernatants with Ni-NTA resin were loaded on a gravity flow column and washed with the PBS buffer. The proteins were eluted in the PBS buffer additionally containing 200 mM imidazole, 30 mM NaCl and 5 mM Tris-HCl, pH 8.0. All of the following steps were performed under red light to avoid photodamage. The retinal was added to the eluted proteins in approximately two-fold molar excess. Resulting mixtures were dialysed overnight against the PBS buffer. During the dialysis, the color of the samples changed from orange to pinkish red, indicating retinal binding. After the dialysis, samples were again incubated overnight with Ni-NTA resin with constant stirring at 4 °C. Resulting solutions with Ni-NTA resin were loaded on a gravity flow column and washed with the PBS buffer to get rid of excess retinal. The protein was eluted in the PBS buffer additionally containing 200 mM imidazole, 30 mM NaCl and 5 mM Tris-HCl, pH 8.0. Resulting solution contained protein in two stable soluble forms - orange (presumably misfolded) oligomeric state and pink monomeric state. We performed size exclusion chromatography (SEC) using Superdex 75 Increase 10/300 GL column (Cytiva, USA) to separate the two forms. Samples purity was assessed by SDS-PAGE (Fig. S1) and SEC (Fig. 1f,g).

For pH sensitivity analyses and transient absorption spectroscopy, a series of buffers was prepared, each containing 75 mM NaCl and 10 mM of MES, MOPS, HEPES, CAPS, and TRIS. pH values were adjusted by incrementally adding HCl or NaOH until reaching the desirable values. The concentrated protein solution in the PBS buffer was mixed in a 1:7 volume ratio with the prepared buffers; resulting pH value was determined by measuring separately the pH value of a large volume 1:7 mixture of the buffers without the protein.

### Thermal stability

For determination of thermal stability of NeuroBR_A,C, protein solutions in the PBS buffer were heated up from 10 to 85 °C at 1 °C/minute rate. Absorption spectra were recorded using AvaSpec-2048L spectrometer (Avantes). Samples were illuminated by AvaLight-DHc full-range light source (Avantes). The proportion of the orange misfolded form was determined based on the ratio of the absorption peaks observed at 523 and 359 nm.

### Dark adaptation

NeuroBR_A and C in PBS buffer pH 7.5 were incubated overnight and transferred to the cuvette holder in complete darkness at 4 °C. Absorption spectra were measured as described above. After the measurement, the samples were intensely illuminated and returned back to the cuvette holder for subsequent measurement.

### Transient absorption spectroscopy

The transient absorption changes were measured using a custom-built setup^37^. In brief, protein activation was achieved with a Brilliant BRILL/IR-10 Nd:YAG laser coupled with a Rainbow OPO (420-680 nm, Quantel, France), delivering near-nanosecond pulses at 530 nm wavelength with an energy of approximately 1 mJ per pulse. A 75 W Xe-arc lamp (Hamamatsu, Japan) served as the probing light source. The samples were positioned between two synchronized monochromators (LSH-150, LOT, Germany) controlled by stepper motors. A photomultiplier tube (R12829, Hamamatsu Photonics, Japan) detected the light. Data were collected from 330 to 730 nm with a 10 nm step size, with each wavelength measurement averaged over 10 repetitions. The sample temperature was maintained at 20°C using a temperature-controlled cuvette holder. Data acquisition was carried out using two digital oscilloscopes (Keysight DSO-X 4022A) in overlapping time windows.

Transient absorption changes were fitted with multiexponential global fit with MEXFIT^38^. Both for NeuroBR_A and NeuroBR_C, 3 time constants were used. After that, the model of irreversible sequential transitions was used to obtain absorption spectra of intermediates^39^. Absorption spectra of intermediates and ground states were fitted with skewed Gaussian functions^39^ for absorption peaks and a power function with a negative exponent for background scattering.

### Crystallization, data collection, and structure determination

The monomeric fractions of NeuroBR_A,C were concentrated and crystallized by a sitting drop vapor diffusion approach using the NT8 robotic system (Formulatrix, USA). The drops contained 150 nL concentrated protein solution and 150 nL reservoir solution. Crystallization plates were stored at 20 °C in the dark. The rectangular crystals of NeuroBR_A appeared after 2 days in the probes containing 0.2 M Ammonium sulfate and 30 % w/v PEG 4000 or PEG 8000 as precipitant solutions. The crystals were harvested using micromounts under red light illumination, cryoprotected by immersing into 15 % glycerol, flash-cooled and stored in liquid nitrogen. Diffraction data were collected at the BL02U1 beamline of the Shanghai Synchrotron Radiation Facility (SSRF). Data integration was performed using XIA2 software^40–44^, and scaling was completed with the STARANISO web server^45^, revealing anisotropic diffraction limits of 1.76, 1.78, and 2.21 Å (Table S2). Elliptically truncated data were subsequently utilized for molecular replacement in Phaser^46^, using the AF2-predicted model as the template. Structure refinement was conducted in Refmac5^47,48^, with related statistics provided in Table S3.

## Data availability

Molecular dynamics trajectories were deposited to Zenodo and are available using the following link: https://zenodo.org/doi/10.5281/zenodo.14194414. Crystallographic structure factors and coordinates for NeuroBR_A were deposited into the Protein Data Bank (PDB) under accession code 9KME.

## Acknowledgements

Computational parts of the work were supported by the Ministry of Science and Higher Education of the Russian Federation, agreement 075-03-2024-117 (project FSMG-2021-0002, to I.Yu.G.). Functional studies were supported by the Russian Science Foundation (21-64-00018, to V.B.). Synchrotron data collection was supported by the Ministry of Science and Higher Education of the Russian Federation (grant no. 075-15-2021-1354, to I.K.). We thank SSRF for providing the opportunity to collect crystallographic data at the beamline BL02U1.

## Author Contributions

I.G. and A.N. conceived the project. I.G. supervised the project. A.N. developed the computational pipeline and obtained NeuroBR sequences. P.S. and A.K. conducted molecular dynamics simulations. A.A. prepared the phylogenetic tree. A.N. expressed, purified and characterized the proteins. Y.A., Y.S.G., E.K., A.M. and O.S. helped with experiments. F.T. measured the photocycle data. I.C. supervised the photocycle data processing. A.R. obtained the crystals. V.B. and I.K. collected and processed the diffraction data. I.G. and A.N. solved and refined the structure. A.N. and I.G. prepared the original draft of the manuscript with contributions from all coauthors. All authors contributed to the preparation of the final version and reviewed it.

