## Supplementary Information for "Engineering of soluble bacteriorhodopsin"

**Supplementary Text**

**Data S1-S2**

**Tables S1-S3**

**Figures S1-S5**

### Supplementary Text

#### Characterization of NeuroBR_B

Reconstituted holo-form of NeuroBR_B exhibits extremely low thermal- and photo- stability. Judged by the ratio of orange (absorbing at ~400 nm, presumably with deprotonated retinal Schiff base) and pink (absorbing at ~530 nm, presumably with protonated retinal Schiff base) forms, the protein was not fully folded at any temperature, with the maximum amount (~50%) of pink form observed between 20 and 30 °C, both in the PBS buffer, pH = 7.5, and in the low salt buffer containing 10 mM NaPi, pH = 8. Increasing or decreasing pH from the range of 7 to 8 causes the protein to transition to orange form. Although no quantitative measurements were made, we note that NeuroBR_B had significantly weaker photo-stability compared to NeuroBR_A and C, quickly converting into the orange form under daylight. Being unable to obtain a stable pink form of NeuroBR_B and having much better variants A and C, we did not pursue further characterization.

**Data S1**. Amino acid sequences of engineered soluble BR variants. Structural protein amino acids are shown as capital letters and extra residues (flexible linkers and hexahistidine tags) are shown as small letters. The hexahistidine tags used for metal-affinity purification are underlined.

>NeuroBR_A

MSMRPESKYYEKATEEMEKAYEEFKKKGEGVEDEEAKKFYDLLTEVPRIAYEQYKKILDGEGIEKVEVDGKEIEVPVAR
YEDWEKTTPLLLEVLANLVDASEELKEKLISKAKEMITLGKKGALETDPEKRFEYWKKSTEKMNEIIDLLENGFKENLS
SLKPERKKTYEEARKLTIELWSKYPEIWKKGPLGEGKVPLEETVKQFTELDVSAKVGFGELVLSSEAIYSgsghhhhhh

>NeuroBR_B

MSSRPESEAYAAATKEMTDAAAKFKEKGKGEKDPEAQKFYELLTKVPAIAAESYQALLDGSGLVRVRVDGKDVEVPVAR
YDDWAVTTPLLLEVLALLVNASEELKQKLLSLAQQMIDLGRQGALETDPEKRFEYWDKSTAAMNEIFDLLENGFEENLS
SLKPERLETYNKLRKMTLELWSQYPEIWRRGPLGKGEVPLAETAARFRELDVSAKVGFGEIVTSSKAIYSgsghhhhhh

>NeuroBR_C

MADRPEAAALAAATAAMTAAAAAFAAKGAGETDPEAQKFYELATKVPAIAAASYQAMLDGSGIVLVEVDGKEVEVYVAR
YDDWKVTTPLLLEILALLVEAKEEVKKELIELANKMIDLGEKGALETEPEKRFEYWEKSTEYMNKIIDILKNGFEENLE
SLEPERKETFEKLKEMTIKLWSQYPKIWREGTLGEGKVSLEEEVARFAELDVSAKVGFGDLLLSSKAIYSgsghhhhhh

**Data S2**. Nucleotide sequences of BR variants analyzed in this work. Nuclease cleavage sites are underlined. Sequences were designed for cloning into pET-28a(+) plasmid via XbaI (TCTAGA) and BamHI (GGATCC) restriction sites.

>NeuroBR_A
TCTAGAAATAATTTTGTTTAACTTTAAGAAGGAGATATACATATGAGTATGCGTCCTGAGAGCAAATATTACGAAAAAGCAACCGA
AGAGATGGAAAAAGCCTATGAGGAGTTCAAGAAGAAGGGCGAGGGTGTCGAAGATGAAGAGGCCAAAAAGTTTTATGATCTGCTGA
CCGAAGTTCCGCGTATTGCCTATGAACAGTATAAGAAAATTCTGGATGGCGAGGGCATTGAAAAAGTTGAAGTTGATGGCAAAGAA
ATCGAAGTGCCGGTTGCACGTTATGAAGATTGGGAGAAAACCACACCGCTGCTGCTGGAAGTTCTGGCAAATCTGGTTGATGCAAG
CGAAGAACTGAAAGAGAAACTGATTAGCAAAGCCAAAGAGATGATTACTTTAGGTAAAAAGGGTGCACTGGAAACCGATCCTGAAA
AACGCTTTGAATATTGGAAGAAAAGCACCGAGAAAATGAACGAGATTATTGACTTGCTGGAAAACGGCTTTAAAGAAAATCTGAGC
AGCTTGAAGCCCGAACGTAAGAAAACGTATGAAGAAGCACGTAAACTGACCATTGAACTGTGGTCAAAATACCCGGAAATTTGGAA
AAAGGGTCCGTTAGGTGAAGGCAAAGTTCCGCTGGAAGAAACCGTTAAACAGTTTACCGAACTGGATGTTAGCGCCAAAGTTGGTT
TTGGTGAACTGGTCCTGAGCAGTGAAGCAATTTATAGCGGTAGCGGTCATCATCACCATCATCATTAAGGATCC

>NeuroBR_B
TCTAGAAATAATTTTGTTTAACTTTAAGAAGGAGATATACATATGAGCAGCCGTCCTGAAAGCGAAGCCTATGCAGCAGCAACCAA
AGAAATGACCGATGCAGCCGCTAAATTCAAGGAAAAAGGTAAAGGCGAAAAAGACCCGGAAGCACAGAAATTCTATGAACTGCTGA
CCAAAGTTCCGGCTATTGCAGCAGAAAGCTATCAGGCACTGCTGGATGGTAGCGGTCTGGTTCGTGTTCGTGTTGATGGTAAAGAT
GTTGAAGTTCCGGTTGCACGTTATGATGATTGGGCAGTTACCACACCGCTGCTGCTGGAAGTTCTGGCCCTGCTGGTTAATGCAAG
CGAAGAACTGAAACAGAAACTGCTGAGTCTGGCACAGCAGATGATTGATCTGGGTCGTCAGGGTGCACTGGAAACCGATCCTGAAA
AACGCTTTGAATATTGGGATAAAAGCACCGCAGCCATGAACGAAATTTTTGATCTGCTGGAAAATGGCTTCGAAGAAAATCTGAGC
AGCCTGAAACCGGAACGTCTGGAAACCTATAACAAACTGCGTAAAATGACCCTGGAACTGTGGTCCCAGTATCCTGAAATTTGGCG
TCGTGGTCCGTTAGGTAAAGGTGAAGTGCCGCTGGCAGAAACCGCAGCACGTTTTCGTGAACTGGATGTTAGCGCAAAAGTTGGTT
TTGGTGAAATTGTTACCAGCAGCAAAGCAATTTATAGCGGTAGTGGTCATCATCATCACCATCACTAAGGATCC

>NeuroBR_C
TCTAGAAATAATTTTGTTTAACTTTAAGAAGGAGATATACATATGGCAGATCGTCCTGAAGCTGCGGCACTGGCCGCGGCAACCGC
CGCAATGACTGCTGCTGCAGCAGCATTTGCAGCCAAAGGTGCCGGTGAAACCGATCCTGAAGCTCAGAAATTCTATGAACTGGCAA
CCAAAGTTCCGGCTATTGCAGCAGCCAGCTATCAGGCAATGCTGGATGGTAGCGGTATTGTTCTGGTTGAAGTTGATGGTAAAGAG
GTCGAAGTTTACGTTGCACGTTATGATGATTGGAAAGTTACCACACCGCTGCTGCTGGAAATTCTGGCACTGCTGGTAGAAGCCAA
AGAAGAAGTTAAGAAAGAACTGATTGAGCTGGCCAACAAAATGATTGATCTGGGTGAAAAAGGTGCACTGGAAACCGAACCGGAAA
AACGCTTTGAATATTGGGAAAAATCCACCGAGTACATGAACAAGATCATCGACATTCTGAAAAACGGCTTCGAAGAAAATCTGGAA
AGCCTGGAGCCCGAACGTAAGGAAACCTTTGAAAAACTGAAAGAAATGACCATCAAGCTGTGGTCACAGTATCCGAAAATTTGGCG
TGAAGGTACATTAGGTGAAGGTAAAGTTAGCCTGGAAGAAGAGGTTGCCCGTTTTGCAGAACTGGATGTTAGCGCAAAAGTTGGTT
TTGGTGATCTGCTGCTGAGTAGCAAAGCAATTTATAGCGGTAGCGGTCATCATCACCATCATCATTAAGGATCC

**Table S1**. Parameters of BR variants studied in this work.

| | pLDDT | Alpha Fold model RMSD to 7Z09 (Å) | Number of residues (w/o tag) | Molecular weight (with tag, kDa) | Molecular weight (w/o tag) | pI without tag | $\varepsilon_{280}$ ($M^{-1}$ $cm^{-1}$) | $\varepsilon_{280}$ (mg/mL cm) |
|---|---|---|---|---|---|---|---|---|
| Wild type | 97.0 | 0.90 | 248 | - | 26.8 | - | - | - |
| NeuroBR_A | 96.7 | 0.62 | 227 | 27.5 | 26.5 | 5.15 | 38390 | 0.715 |
| NeuroBR_B | 96.5 | 0.63 | 227 | 26.6 | 25.6 | 5.12 | 33920 | 0.783 |
| NeuroBR_C | 96.3 | 0.60 | 227 | 26.3 | 25.2 | 4.67 | 33920 | 0.774 |

**Table S2:** Diffraction anisotropy information.

Diffraction limits (Å) and corresponding principal axes of the ellipsoid fitted to the diffraction cut-off surface as direction cosines in the orthogonal basis (standard PDB convention), and in terms of reciprocal unit-cell vectors.

| | Diffraction limits (Å) | Principal axes in the orthogonal basis | Principal axes reciprocal unit-cell vectors |
|---|---|---|---|
| Diffraction limit #1 | 1.755 | ( 1.0000, 0.0000, 0.0000) | a* |
| Diffraction limit #2 | 1.778 | ( 0.0000, 1.0000, 0.0000) | b* |
| Diffraction limit #3 | 2.212 | ( 0.0000, 0.0000, 1.0000) | c* |

Eigenvalues of the overall anisotropy tensor on |F|s (Å$^2$), the same eigenvalues after subtraction of the smallest eigenvalue (as used in the anisotropy correction), and corresponding eigenvectors of the overall anisotropy tensor as direction cosines in the orthogonal basis (standard PDB convention), and in terms of reciprocal unit-cell vectors:

| | Eigenvalues (Å$^2$) | Eigenvalues used in the anisotropy correction (Å$^2$) | Eigenvectors in the orthogonal basis | Eigenvectors in reciprocal unit-cell vectors |
|---|---|---|---|---|
| Eigenvalue #1 | 13.24 | 0.00 | ( 1.0000, 0.0000, 0.0000) | a* |
| Eigenvalue #2 | 16.92 | 3.68 | ( 0.0000, 1.0000, 0.0000) | b* |
| Eigenvalue #3 | 33.57 | 20.33 | ( 0.0000, 0.0000, 1.0000) | c* |

**Table S3**. Crystallographic data collection and refinement statistics.

| | |
|---|---|
| PDB ID | 9KME |
| Wavelength, Å | 0.979 |
| Resolution range*, Å | 36.17 - 1.76 (1.88 - 1.76) |
| Space group | P 21 21 21 |
| Unit cell, Å ° | 95.03 96.09 109.93 90 90 90 |
| Total reflections | 980346 (39576) |
| Unique reflections | 76128 (3849) |
| Multiplicity | 12.9 (10.3) |
| Completeness (spherical), % | 76.7 (22.2) |
| Completeness (ellipsoidal), % | 95.8 (70.2) |
| Mean I/σ(I) | 15.2 (2.5) |
| Wilson B-factor*, Å$^2$ | 16.5 |
| R-merge | 0.127 (0.968) |
| R-meas | 0.132 (1.019) |
| R-pim | 0.036 (0.310) |
| CC1/2 | 0.999 (0.655) |
| Resolution range used in the refinement, Å | 36.17 - 1.76, 1.78, 2.21 |
| Reflections used in refinement | 70884 |
| Reflections used for R-free | 236 |
| R-work | 0.1771 |
| R-free | 0.2125 |
| Number of non-hydrogen atoms | 7944 |
| macromolecules | 7423 |
| ligands | 20 |
| solvent | 501 |
| Protein residues | 903 |
| $RMS_{bonds}$, Å | 0.008 |
| $RMS_{angles}$, ° | 1.18 |
| Ramachandran favored, % | 99 |
| Ramachandran allowed, % | 1 |
| Ramachandran outliers, % | 0 |
| Rotamer outliers, % | 1 |
| Clashscore | 11 |
| Average B-factor, Å$^2$ | 22.8 |
| macromolecules | 22.4 |
| ligands | 45.6 |
| solvent | 28.3 |
| Number of TLS groups | 4 |

Statistics for the highest-resolution shell are shown in parentheses.

* - anisotropy information is presented in Table S2.

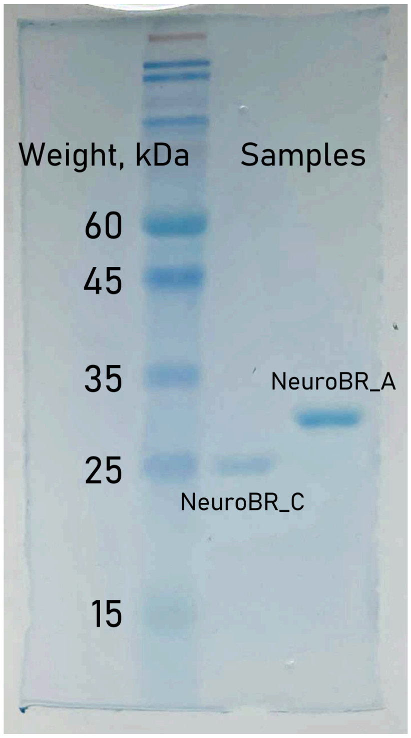


**Figure S1.** SDS-PAGE of NeuroBR_A and C samples purified as described in the Methods section.

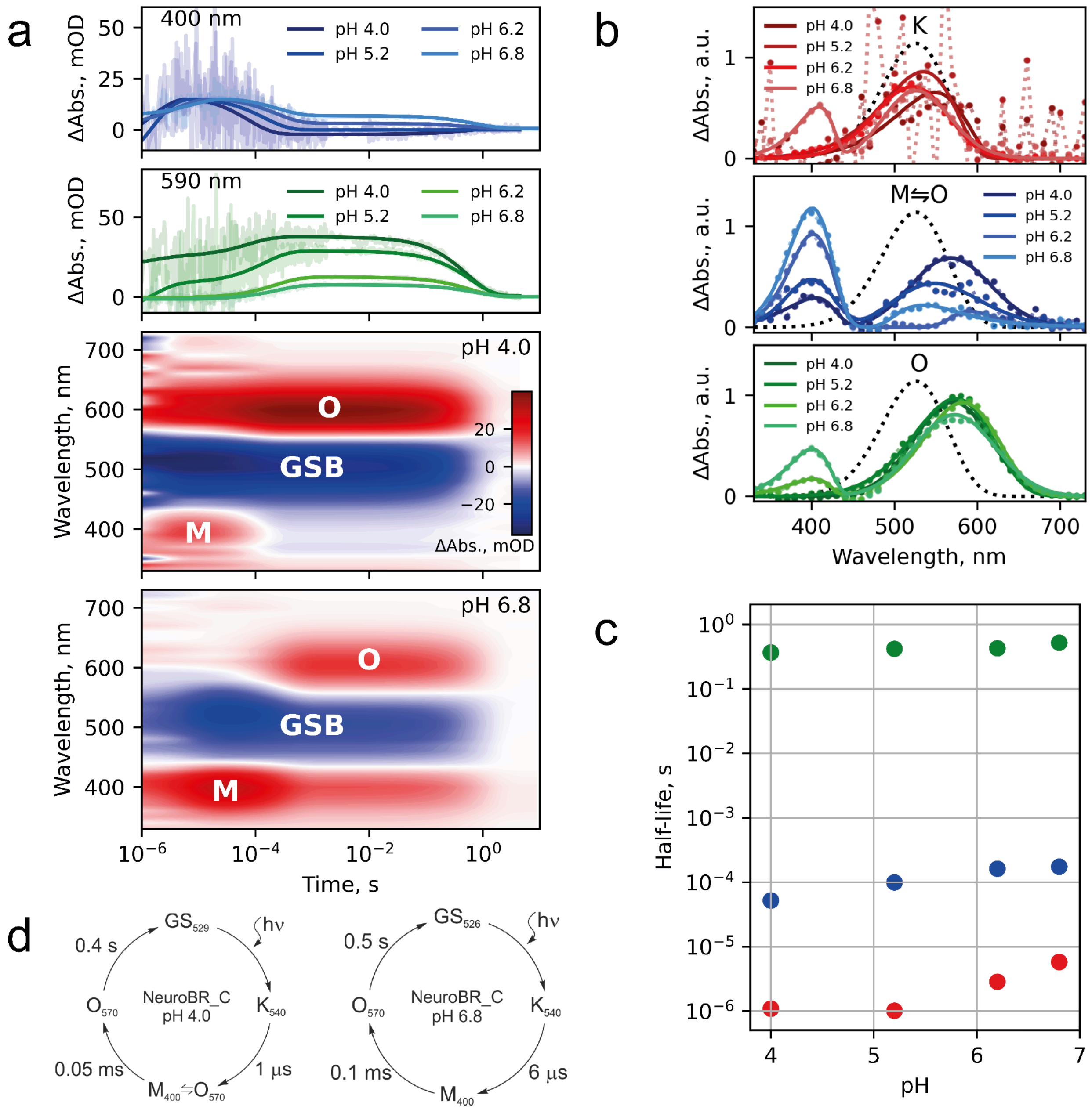


**Figure S2. Transient absorption spectroscopy of NeuroBR_C. a**, Changes in absorbance of NeuroBR_C in solution after flash illumination. M and O correspond to areas where absorbance raises due to formation of putative M and O photocycle intermediates. GSB is the ground state bleaching area where absorbance corresponding to the ground state is diminished due to formation of photocycle intermediates. **b**, Recovered absorption spectra of NeuroBR_C photocycle intermediates at different buffer pH values. **c**, Dependence of NeuroBR_C photocycle intermediate half lifes on pH. **d,** Model of NeuroBR_C photocycles at pH 4.0 and pH 6.8.

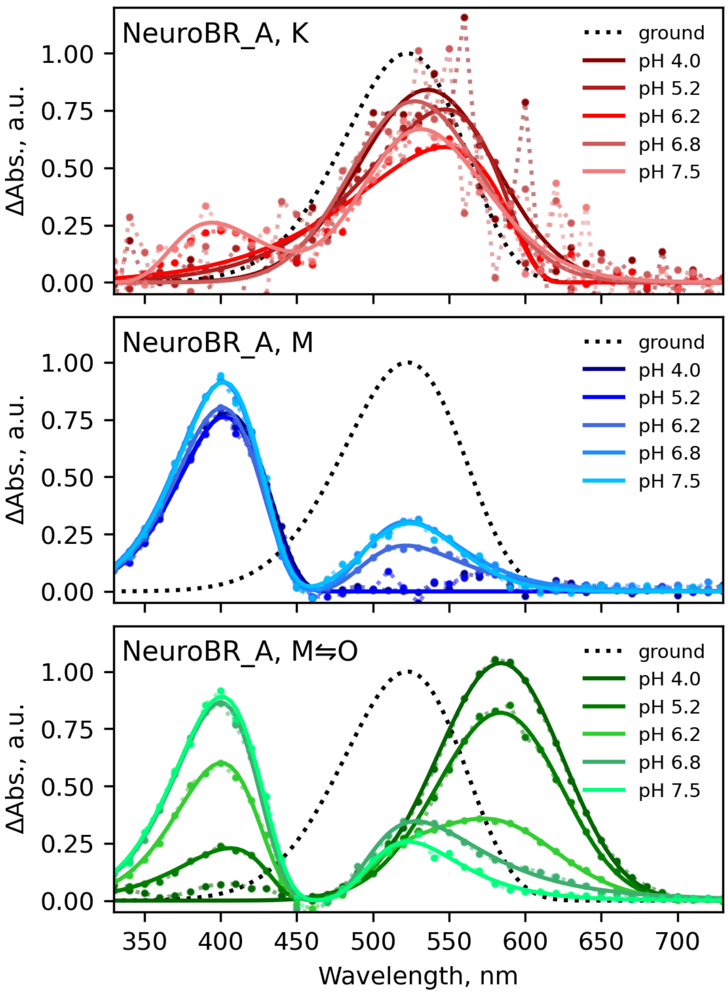


**Figure S3.** Recovered absorption spectra of NeuroBR_A intermediates at different buffer pH values.

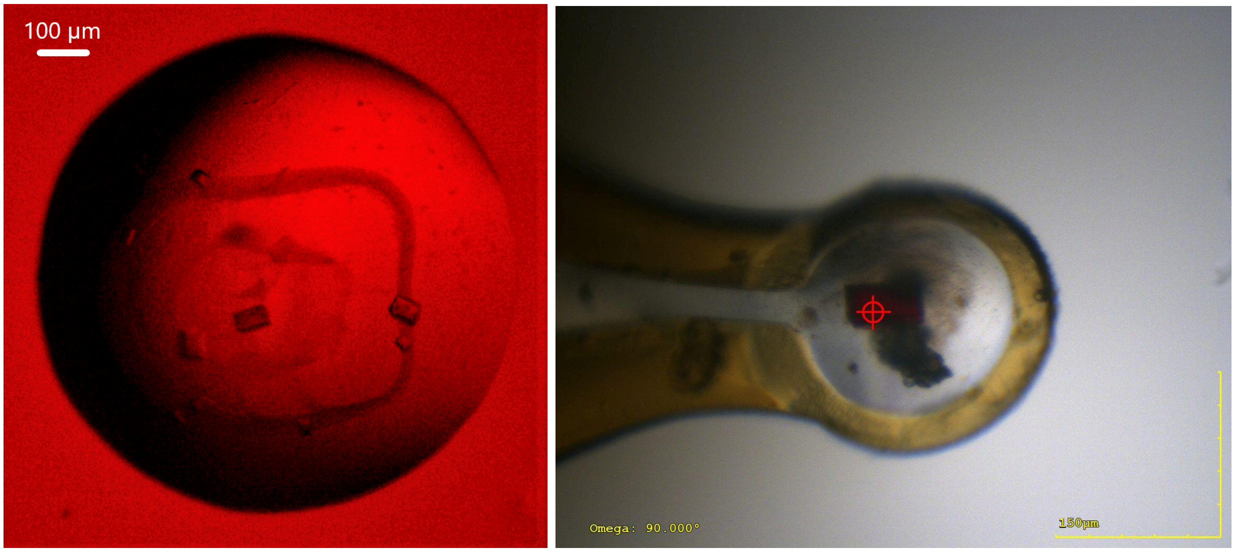


**Figure S4. NeuroBR_A crystals.** (left) Crystals in the crystallization drop observed via a microscope under red light illumination. (right) Mounted crystal imaged during diffraction data collection at SSRF.

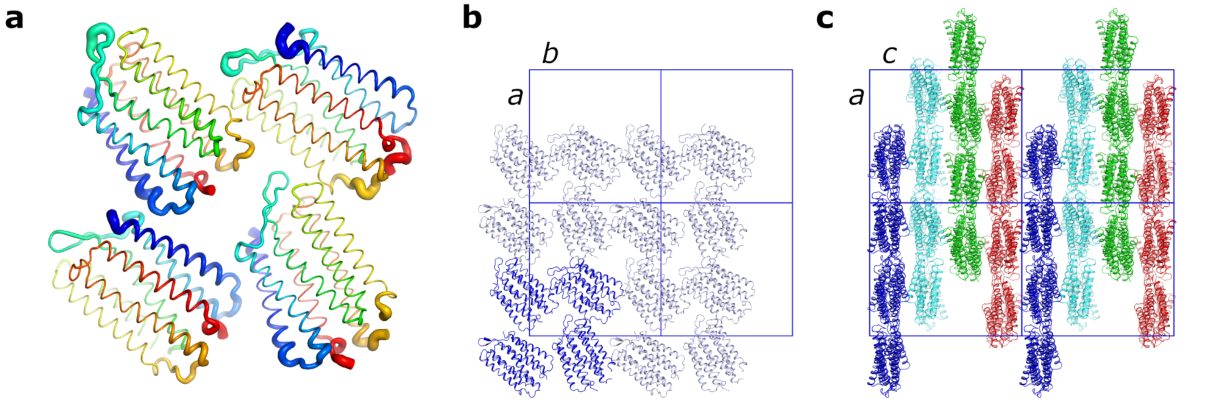


**Figure S5. Crystal packing of NeuroBR_A. a**, Relative distributions of the B-factor values for the four NeuroBR molecules present in the asymmetric unit cell (ASU). Coloring changes from blue to red from the N-terminus to the C-terminus. **b**, Packing in the *a*-*b* plane. ASU contents of one cell is shown in blue and contents of the adjacent cells is shown in light blue. *a* = 95.03 Å, *b* = 96.09 Å. **c**, Packing in the *a*-*c* plane. Each cell contains 16 copies of NeuroBR, organized as four parallel layers of four NeuroBR molecules. Crystallographic symmetry-related molecules are colored blue, light blue, green and red. *a* = 95.03 Å, *c* = 109.93 Å.
